# Co-encapsulated liposomal formulations of tubacin and erlotinib.HCl: a comprehensive stability strategy

**DOI:** 10.1101/2025.10.08.681158

**Authors:** Cindy Schelker, Catia Monteiro, Gerrit Borchard, Patrycja Nowak-Sliwinska

## Abstract

The co-encapsulation of two anticancer drugs within liposomes is a promising strategy for delivering a synergistic drug ratio to tumors. In this preliminary study, we systematically investigated the formulation stability of liposomes co-encapsulating tubacin and erlotinib.HCl, as well as the development of a freeze-drying (FD) method as a stabilization strategy. Notably, the effect of cryoprotectants on size and drug encapsulation conservation was studied both before and after freezing and thawing (FD).

Tubacin was passively loaded into the lipid bilayer, while erlotinib.HCl was remotely loaded via an ammonium sulfate gradient. Lipid composition studies revealed that erlotinib.HCl encapsulation was the highest at pH 3. Additionally, DSPC:Chol ratios significantly impacted drug loading: high cholesterol content improved erlotinib.HCl retention but reduced tubacin encapsulation. Despite optimization, substantial drug losses occurred during storage at 4°C, leading to the development of freeze-drying as a stabilization strategy. Sucrose and trehalose were evaluated as cryoprotectants at different concentrations. Sucrose (10% w/v) demonstrated higher cryoprotection, maintaining liposome sizes within a range optimal for the enhanced permeability and retention (EPR) effect. However, the inclusion of internal sucrose did not further improve stability. Process optimization identified 150 mM ammonium sulfate as the optimal concentration for remote loading while preserving formulation stability. Comparative evaluation of two freeze-drying protocols revealed that the controlled stepwise system (Telstar^®^) outperformed the alternative approach, demonstrating superior drug retention and particle size preservation. Asymmetric flow field-flow fractionation coupled with dynamic light scattering and multi-angle light scattering (AF4-DLS-MALS) analysis confirmed minimal particle aggregation post-lyophilization, with preserved monodispersity and spherical morphology.

These findings highlight the critical importance of cryoprotectant selection, buffer optimization, and freeze-drying protocol design for developing stable, co-encapsulated liposomal formulations.

## Introduction

In the fight against cancer, chemotherapy has been the standard of care for a long time. The use of drug combination therapies demonstrated superior clinical benefits compared to a single active pharmaceutical ingredient (API), primarily by overcoming the rapid drug resistance development observed with monotherapies [1]. Classical API dose selection for combination therapies is typically based on a drug’s maximum tolerated dose (MTD) to maximize therapeutic efficacy. However, this traditional approach presents critical limitations, i.e. it leads to dose-limiting toxicity [2] and it fails to account for potential drug-drug interactions that could potentiate therapeutic outcomes. Compared to classic drug combinations, synergy-based API combinations enable drug dose reduction while enhancing target selectivity and minimizing resistance occurrence [3, 4]. These benefits arise from maintaining precise ratios between combined APIs. Consequently, a major translational challenge persists to ensure tumor exposure to the synergistic API ratios, primarily due to divergent pharmacokinetic profiles of the individual APIs.

Liposomes represent the most successful type of drug nanocarriers on the global market [5]. This can be explained by the liposome architecture with an aqueous core enclosed by a phospholipid bilayer, allowing the encapsulation of APIs of various log P values [6]. Co-encapsulation of APIs within liposomes is a common strategy for drug co-delivery and maintenance of the API ratio *in vivo* [7, 8]. They are versatile and potentially allow for an accumulation at the tumor site [9]. Tolcher and Mayer initiated the encapsulation of synergistic drug ratios within liposomes, eventually leading to the development of Vyxeos^®^ [2]. To date, no other liposomal formulation co-encapsulating two APIs has received regulatory approval. The currently marketed version exists as a lyophilized formulation [10], which raises an additional consideration regarding API retention during storage. Liposomes in solution are prone to instability due to phospholipid hydrolysis and chemical degradation [11, 12]. While freeze-drying (FD) or lyophilization represents an established strategy to enhance storage stability, the process may significantly alter the liposome structure. The addition of cryoprotectants has been shown to mitigate these FD-induced alterations [13].

Disaccharides such as sucrose and trehalose are often described as highly effective cryoprotectants [14]. Their mechanism of action is not fully elucidated, and so far, there are three primary theories proposed. One theory described by Crowe and Crowe [15], suggests that the water is replaced by disaccharides, bridging phospholipids through hydrogen bonds, replacing water molecules during dehydration [16, 17]. This explains why successful cryoprotectants such as disaccharides often display a large number of hydroxyl groups. The second theory is the vitrification or glass formation proposed by Koster *et al*. [18], where the molecular mobility is reduced due to the viscous matrix surrounding the bilayers. Liposomes collapse during freeze-drying when the temperature used during the process exceeds the glass transition temperature (Tg) [19]. The vitrification process depends on the Tg of the cryoprotectants used. At temperatures exceeding the Tg, the viscosity of the phospholipid bilayer decreases while molecular mobility increases [16]. It is accepted that these two theories do not exclude each other but are rather complementary explanations [19]. Finally, the third theory of the kosmotropic effect, where cryoprotectants interact with water, reducing its interaction with the liposomal membrane surfaces [20].

Despite the success of Vyxeos^®^, no study has explored the impact of cryoprotectant selection and freeze-drying processes on liposomes for API co-delivery. Current literature reveals a lack of detailed information regarding lyophilization processes for liposomal formulations. In this context, our study systematically addresses this critical knowledge gap by (i) investigating two distinct freeze-drying protocols, and (ii) comparing the use of trehalose and sucrose as cryoprotectants. This work aimed to establish optimized storage conditions that maintain liposomal stability of liposomal formulations for optimal synergistic API delivery.

## Materials and Methods

### Compounds

Erlotinib.HCl was purchased from LC Labs (Woburn, MA, USA). Tubacin (purity ≥ 98% HPLC), and cholesterol (CHOL), D-α-tocopheryl polyethylene glycol 1000 succinate (TPGS), Triton X-100^®^, anhydrous DMSO, phosphate salt (H_2_KO_4_P), ammonium sulfate (AS) and sucrose were all purchased from Sigma Aldrich Chemie GmbH (Schnelldorf, Germany). 1,2-dipalmitoylphosphatidylcholine (DPPC) and 1,2-distearoyl-sn-glycero-3-phosphatidylcholine (DSPC) were purchased from Avanti Research (Birmingham, AL, USA). Methanol, acetonitrile, and ethanol were obtained from Fischer Chemical (Reinach, Switzerland). Trehalose dihydrate ≥ 99% was obtained from, Fluka Chemie GmbH (Buchs, Switzerland). Citric acid monohydrate was purchased from Hänseler AG (Herisau, Switzerland). Sephadex G-25 Hitrap desalting column was purchased from Cytiva (Rosersberg, Sweden), electronic pipette was from Eppendorf (Hamburg, Germany).

### UPLC quantification

The chromatographic separation of erlotinib.HCl was done using a Waters Acquity System (Milford, MA, USA) equipped with a binary solvent delivery pump, autosampler, sample manager, and a photodiode array (PDA) detector. Separation was carried out on an Acquity UPLC^®^ BEH C18 2.1 × 100 mm, 1.7 µm column. The column was heated to 30°C. The mobile phase was composed of solvent A (phosphate buffer: H_2_KO_4_P 0.05M, pH 6.5 adjusted with NaOH) and solvent B (methanol: acetonitrile 70:30). A gradient method was used starting from a mixture of 50% of A and 50% of B to 100% of B in 6 min. From 6 min up to 6.1 min the mobile phase was a mixture of 50% A and 50% B and stayed at this composition for 7 min. The flow rate was 0.3 ml/min, and UV detection was achieved at 246 nm. The injection volume was 10 µl. Stock solutions of erlotinib.HCl (10 mg/ml) and tubacin (10 mg/ml) were prepared by dissolving drugs in 100% anhydrous DMSO and stored at −20°C until further use. Calibration solutions of erlotinib.HCl and tubacin were prepared under the same conditions as the samples.

### Calibration curve

A standard stock solution of erlotinib.HCl and tubacin in anhydrous DMSO at a concentration of 10 mg/ml was diluted with PBS/Tween 80 (0.1%) at pH 7.4. The final range of concentrations were 0.01, 0.02, 0.05, 0.10, 0.2, and 1 µg/ml. The calibration curve of erlotinib.HCl and tubacin were plotted against the area under the peak on the Y axis and erlotinib.HCl or tubacin concentration on the X axis.

### Liposomes preparation

DSPC:CHOL:TPGS at 8:20:3 molar ratio, liposomes were prepared using the ethanol injection method. For co-loading, tubacin dissolved in DMSO and lipids were dissolved in 1mL of ethanol and were rapidly injected with an electronic pipette into 4 mL of ammonium sulfate (AS) buffer at pH 3. The mixture of ethanol + AS buffer was heated to 60°C for 5 min. Then the mixture was put on ice to cool down. AS buffer was exchanged for citrate buffer 0.1 M at pH 3/sucrose 10%, by size exclusion chromatography (SEC). Buffer exchange by SEC was done by adding 4.5 mL of liposome suspension to the Sephadex column, and 1 mL was eluted with sucrose/citrate buffer and discarded. Then 4.5 mL was eluted with sucrose/citrate buffer and recovered as the purified liposome suspension.

Liposome suspensions were then returned to heat at 60°C in balloons and erlotinib.HCl dissolved in DMSO was added. The mixture was left under stirring for one hour and then cooled on ice. A second SEC was performed to exchange citrate/sucrose buffer to PBS without calcium or magnesium at pH 7.4 and remove free erlotinib.HCl. The SEC protocol used the same volumes as the previous buffer exchange. Single-loaded liposomes with either of the drugs were prepared identically without the drug addition. Liposomal suspensions were stored at 4°C until further use.

### Freeze-drying

The study was conducted using two freeze-dryers to compare their performance. The Christ^®^ freeze-dryer protocol is detailed in **Table 1**, while the Telstar^®^ protocol is detailed in **Table 2**.

**Table 1.** Christ^®^ freeze-dryer protocol.

| **Process** | **Temperature (°C)** | **Pressure (mbar)** | **Duration (hh:mm)** |
| --- | --- | --- | --- |
| Freezing | -80 |  | 01:00 |
| Primary drying | -40 | 0.3 | 24:00 |

**Table 2.** Telstar^®^ freeze-dryer protocol.

| **Process** | **Temperature (°C)** | **Pressure (mbar)** | **Duration (hh:mm)** |
| --- | --- | --- | --- |
| Freezing | -40 |  | 03:00 |
| Condenser preparation |  |  | 00:50 |
| Chamber vacuum |  | 0.3 |  |
| Primary drying | -40 | 0.3 | 25:00 |
| Primary drying | -20 | 0.3 | 05:00 |
| Primary drying | -10 | 0.3 | 03:00 |
| Secondary drying | 22 |  | 04:00 |

### Cryoprotectants

50% (w/v) stock solutions of cryoprotectants sucrose or trehalose were prepared by dissolving 50 g of cryoprotectants in 100 mL water and were then added depending on their final concentration to liposomes at 1 mL. All cryoprotectant concentrations are expressed as %w/v.

### Dynamic light scattering

Mean hydrodynamic size (Z-average) and polydispersity index (PDI) were determined by dynamic light scattering using a Zetasizer (NanoZS, Malvern Panalytical, Malvern, UK) in batch mode. Samples were measured at 25°C in disposable polystyrene cuvettes. The measurement angle was 173°C, the refractive index was set at 1.345, and the absorption at 0.010. The laser attenuator was adjusted automatically. Measurements were performed in triplicate. The equilibration time between samples was 120 seconds. Size and PDI values were measured after dilution at a ratio of 1:20 with PBS 0.9% filtered through filters of 0.22 µm pore size. Data were collected using Zetasizer software v7.13.

### Asymmetrical Flow Field Flow Fractionation (AF4) and multiangle light scattering connected to online DLS in flow-mode

Measurements were conducted by AF4 (AF2000 system, Postnova Analytics, Landsberg, Germany). The system consisted of an autosampler (PN5300), with a solvent organizer (PN7140), degasser (PN7520), and smart stream splitter (PN1650), a FOC pump (PN1130) and a TIP pump (PN1130). The separation channel was equipped with a 10 kDa regenerated cellulose membrane with a 350 µm spacer. The system was coupled to 4 online detectors, a refractive index detector (PN3150), a Multi-Angle Light Scattering (MALS) detector (PN3609), UV/Vis (Waters 2487) detector measuring at a wavelength of λ = 246 nm, and the DLS system (NanoZS, Malvern Panalytical, Malvern, UK). The mobile phase was the same as the external liposome buffer (NaCl 0.9% filtered through 0.1 µm pore size filters). For flow-mode DLS analysis, measurements were obtained with a quartz flow cell (ZEN0023, Malvern Panalytical, Malvern, UK). Gyration radius analysis by the MALS detector was performed using a sphere fit model. The detector flow rate was 0.5 ml/min. The focus step delay time was 3 min; injection flow was 0.20 ml/min with an injection time of 7 min. The crossflow was 1 ml/min, and the focus pump was at 1.30 min. Elution parameters are described in **Table 3**.

**Table 3.** Elution step parameters.

| Step | Time (min) | Cross-flow<br>(ml/min) | Type | Exponent |
| --- | --- | --- | --- | --- |
| 1 | 30 | 1.00 | Linear | 1.0 |
| 2 | 5.0 | 0.19 | Power | 0.8 |
| 3 | 5.0 | 0.10 | Power | 0.8 |
| 4 | 8.0 | 0.06 | Power | 0.8 |
| 5 | 8 | 0.037 | Power | 0.8 |

Finally, the rinse step had a TIP flow of 0.05 ml/min and a focus flow of 0.05 ml/min.

### Drug encapsulation and entrapment efficiency

Erlotinib.HCl and tubacin quantification in liposome formulations was performed using UPLC-UV. 20 µL of the formulation was diluted in 100% DMSO to a final volume of 200 µL in a 2-mL glass vial before determination by the UPLC-UV.

The encapsulation efficiency (EE%) was calculated using Equation 1.

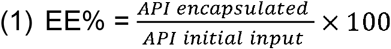

### Statistical analysis

Data analysis was performed using GraphPad Prism software version 10.2.3. A two-way ANOVA was conducted by comparing the data between groups using Tukey’s multiple comparisons test. All *p* values < 0.05 were considered statistically significant.

## Results

DSPC:CHOL:TPGS liposomes at 8:20:3 molar ratio were prepared by the ethanol injection method according to the workflow shown in **Figure 1**.

**Figure 1.**
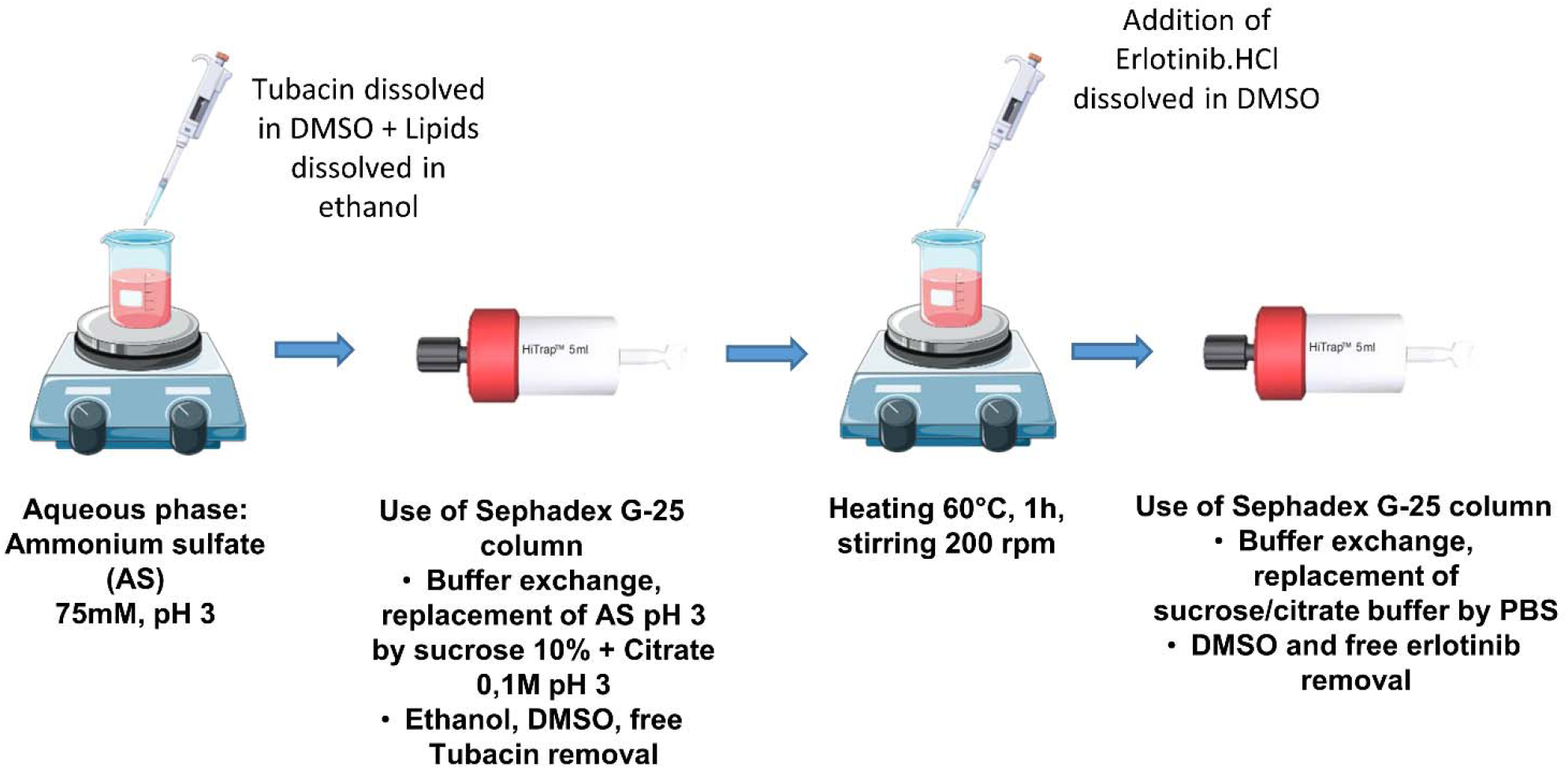
Schematic representation of the workflow for the preparation of co-loaded liposomes.

We investigated the impact of varying internal pH (3-7) values and their impact on drug encapsulation and retention. Results showed that erlotinib·HCl encapsulation exhibited a pH-dependent profile, with optimal loading achieved at pH 3 (85 ± 3% encapsulation efficiency). In contrast, tubacin encapsulation remained unaffected by pH variations (78 ± 4% across all tested pH values). Stability studies at 4°C revealed that after 96 hours, erlotinib·HCl showed consistent leakage rates (25-30%) regardless of pH, while tubacin demonstrated superior retention (<15% loss). These findings established pH 3 as the optimal condition for subsequent studies, as it simultaneously maximized erlotinib·HCl loading while maintaining excellent tubacin encapsulation and satisfactory storage stability for both compounds.

The different pH sensitivity between these drugs highlights the importance of individualized optimization strategies for co-encapsulation systems. (**Figure 2A**). Three DSPC:CHOL molar ratios were tested: 1:3.16, 1:2, and 2:1. The ratio of 1:3.16 led to the highest erlotinib.HCl encapsulation initially at 60% of encapsulation. Molar ratios of 1:2 and 2:1 were less successful, with 50% and 20% of erlotinib.HCl encapsulated, respectively. A molar ratio of 2:1 had the highest tubacin encapsulation, with 60% of drug encapsulation. After one day of storage at 4°C, all liposomal suspensions presented less than 20% of erlotinib.HCl encapsulated. Tubacin retention was higher, with at least 30% of the drug remaining encapsulated after seven days at 4°C (**Figure 2B**). Finally, the impact of single versus co-loading was studied. No impact on the encapsulation efficiencies of either API (**Figure 2B** and **2C**) or on drug loss during storage at 4°C (**Figure 2B** and **2D**) was observed. The DSPC:Chol molar ratio 1:3.16 was selected for the remainder of the study.

**Figure 2.**
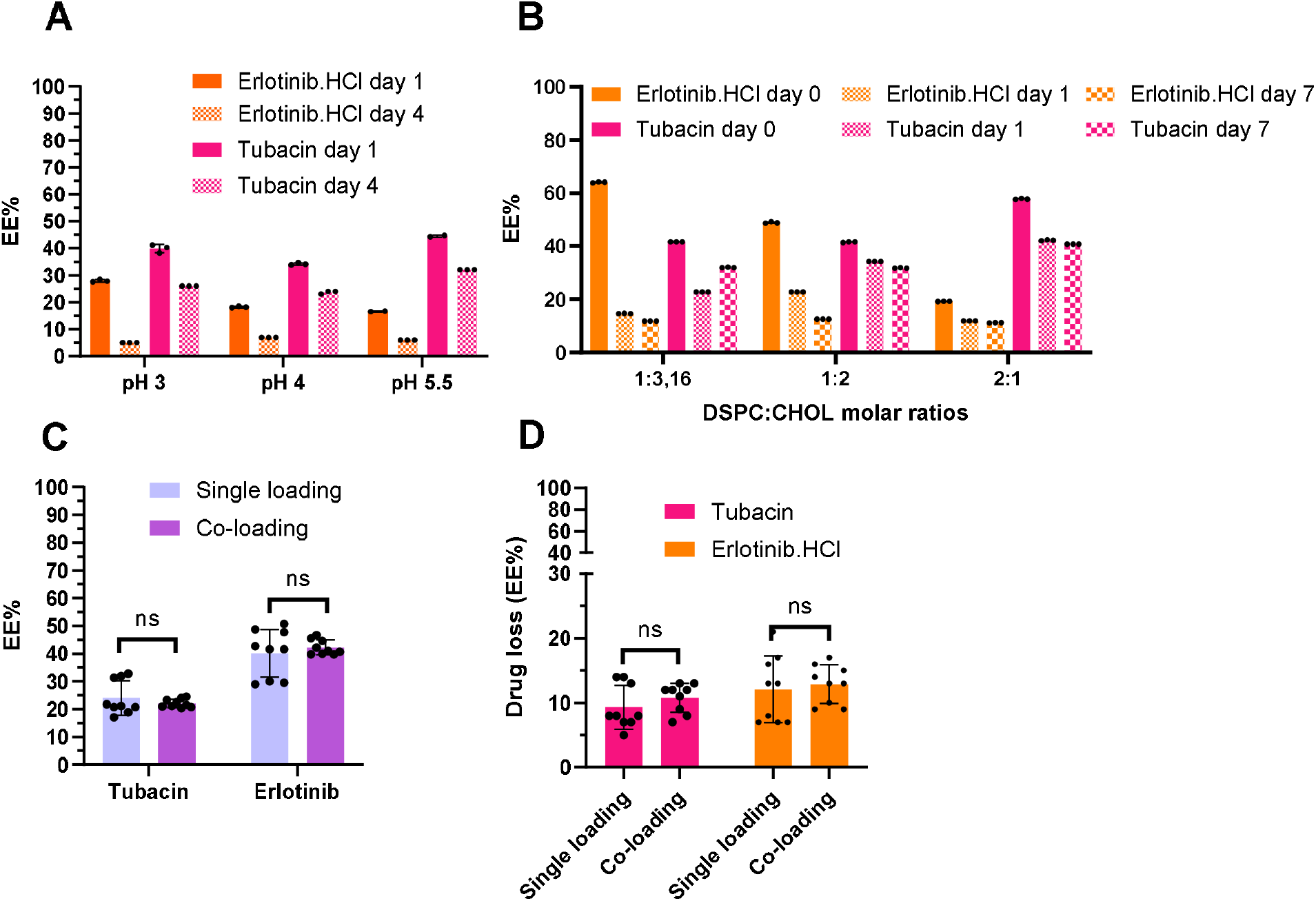
Encapsulation efficiencies and stability during storage at 4°C of co-encapsulated liposomes. Encapsulation efficiencies depend on internal pH (**A**), and on DSPC:CHOL molar ratios (**B**). Impact of single loading versus co-loading on APIs encapsulation efficiencies (**C**) and drug loss during storage at 4°C (**D**). N=1 ± SD (**A** and **B**) and N=3 ± SD (**C** and **D**).

Sizes and PDI of blank liposomes were measured pre- and post-FD. Changes in size and PDI values indicate an insufficient freeze-drying process. Two types of cryoprotectants were incorporated into the formulations before freeze-drying. Before FD, the particle size was around 75 nm with a PDI close to 0.05. Following FD using trehalose at concentrations of 6%, 8%, and 10%, the particle sizes increased to over 250 nm. Sucrose as a cryoprotectant leads to an increase in sizes compared to pre-FD, as well, with sizes reaching 150 nm with sucrose 6%. A trend was observed where higher cryoprotectant concentrations lead to smaller post-FD sizes. Based on these findings, further studies were done with 8% and 10% sucrose (**Figure 3**).

**Figure 3.**
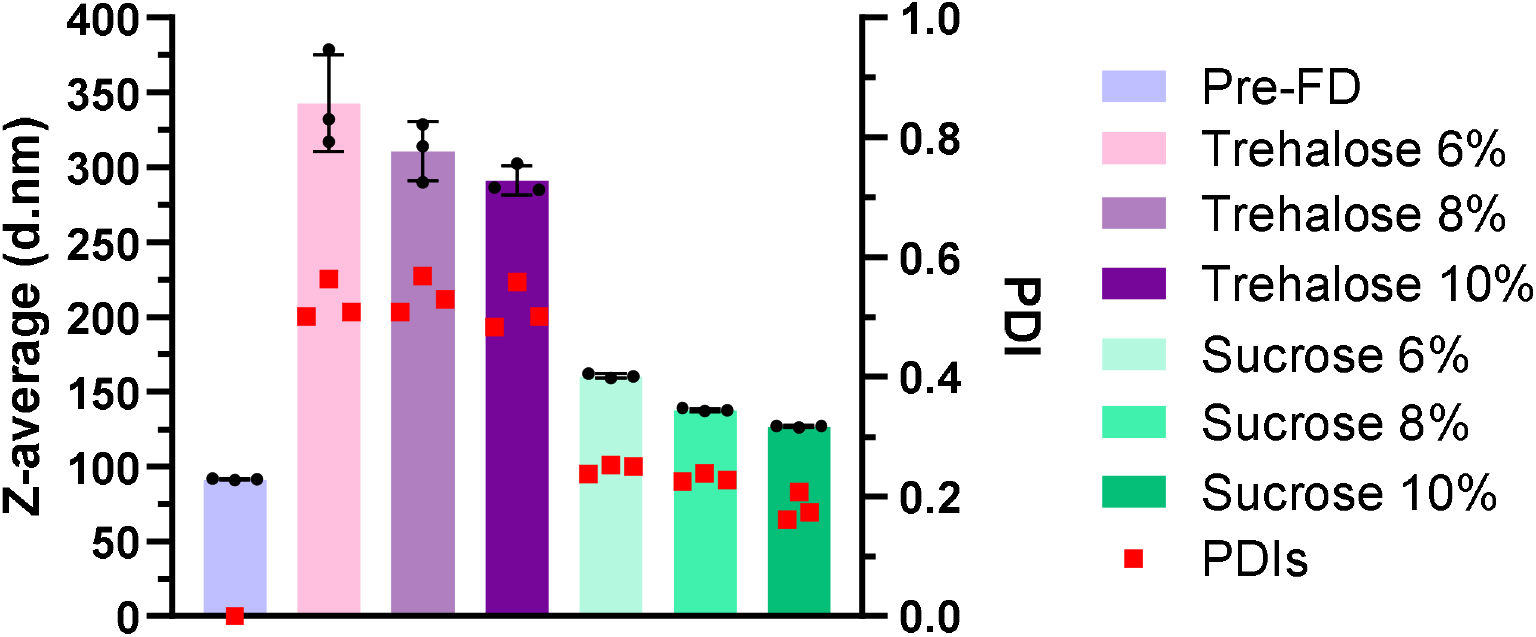
Z-average and PDIs of liposomes pre- and post-FD with different concentrations of trehalose and sucrose. Ammonium sulfate 150 mM, Christ^®^ freeze-dryer (N=1).

Measurement of osmolarities after freeze-drying of LNP post-FD and ammonium sulfate buffer 150 mM was performed and is listed in **Table 4**. The addition of cryoprotectants to liposomes matched the osmolarity of sucrose in the absence of liposomes and was around 600 mmol/kg. Throughout the experimental phase, a standard ammonium sulfate at 150 mM was maintained. yielding a measured osmolarity of 366 mmol/kg. This value represents approximately half that of the osmolarity of cryoprotectants + liposomes. The final external buffer PBS was closer to the value of ammonium sulfate 120 mM, at around 300 mmol/kg.

**Table 4.** Osmolarity of buffers used for the liposomal formulations.

| Buffer | Osmolarity (mmol/kg) |
| --- | --- |
| Blank liposomes + sucrose 8% post-FD | 575.5 ± 5,0 |
| Blank liposomes + sucrose 10% post-FD | 666.0 ± 0 |
| PBS 1x pH 7.4 | 292.0 ± 0 |
| Sucrose 8% in PBS 1x pH 7.4 | 544.5 ± 9.2 |
| Sucrose 10% in PBS 1x pH 7.4 | 657.5 ± 3.5 |
| Ammonium sulfate buffer 120 mM, pH<br>3 | 282.5 |
| Ammonium sulfate buffer 150 mM, pH<br>3 | 366.5 ± 2.1 |
| Ammonium sulfate buffer 300 mM, pH<br>3 | 699.0 ± 8.0 |

(n=2 ± SD)

Initial attempts in the preparation of blank and co-loaded liposomes using 300 mM ammonium sulfate buffer (with either 8 or 10% cryoprotectant) resulted in unstable liposomal formulations, as demonstrated by large particle sizes above 300 nm and PDIs above 0.4 (**Figure 4**).

**Figure 4.**
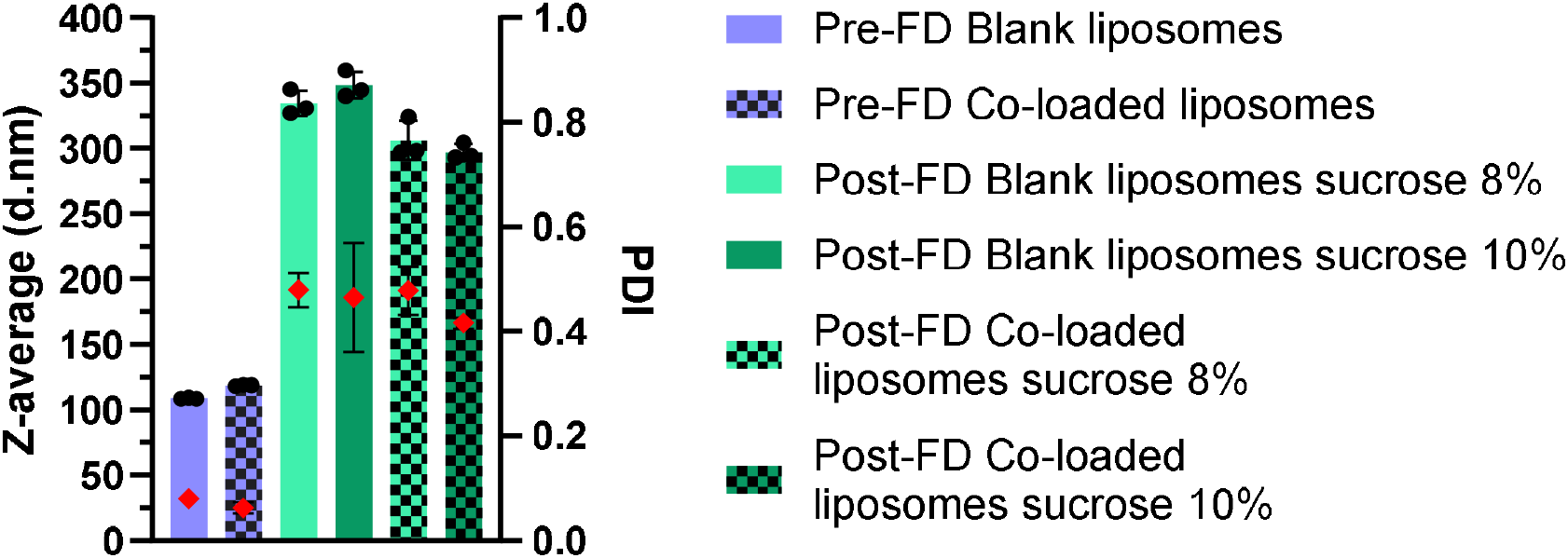
Z-average and PDIs of liposomes pre- and post-FD with different concentrations of sucrose. Ammonium sulfate at 300 mM, Christ^®^ freeze-dryer, (N=1).

Moreover, all formulations, both pre- and post-FD, precipitation within one hour post-formulation was observed. To address these stability challenges, we adjusted the molarity of ammonium sulfate to 120 mM to better match the osmolarity of PBS.

Liposomes formulated with AS at 120 mM exhibited sizes above 600 nm post-lyophilization, increasing from 150 nm to 600 nm (**Figure 5A**). While these formulations showed improved stability with no visible precipitation post-preparation, the elevated PDIs (above 0.5) indicated sub-optimal lyophilization process outcomes. The encapsulation efficiency (EE%) was also assessed. The EE% for tubacin and erlotinib were 90% and 58%, respectively. After freeze-drying, the EE% significantly dropped to 40% and 18% for tubacin and erlotinib, respectively. This degree of API loss was similar to formulations stored at 4°C for 24 hours **(Figure 5B)**. Consequently, we selected AS at 150 mM as the optimal concentration for the co-encapsulation of tubacin and erlotinib.HCl. The lyophilization process did not lead to a difference in tubacin encapsulation, while there was a significant loss of erlotinib.HCl (**Figure 6)**. This selective destabilization suggests distinct molecular interactions between each API and the liposomes during the freeze-drying process.

**Figure 5.**
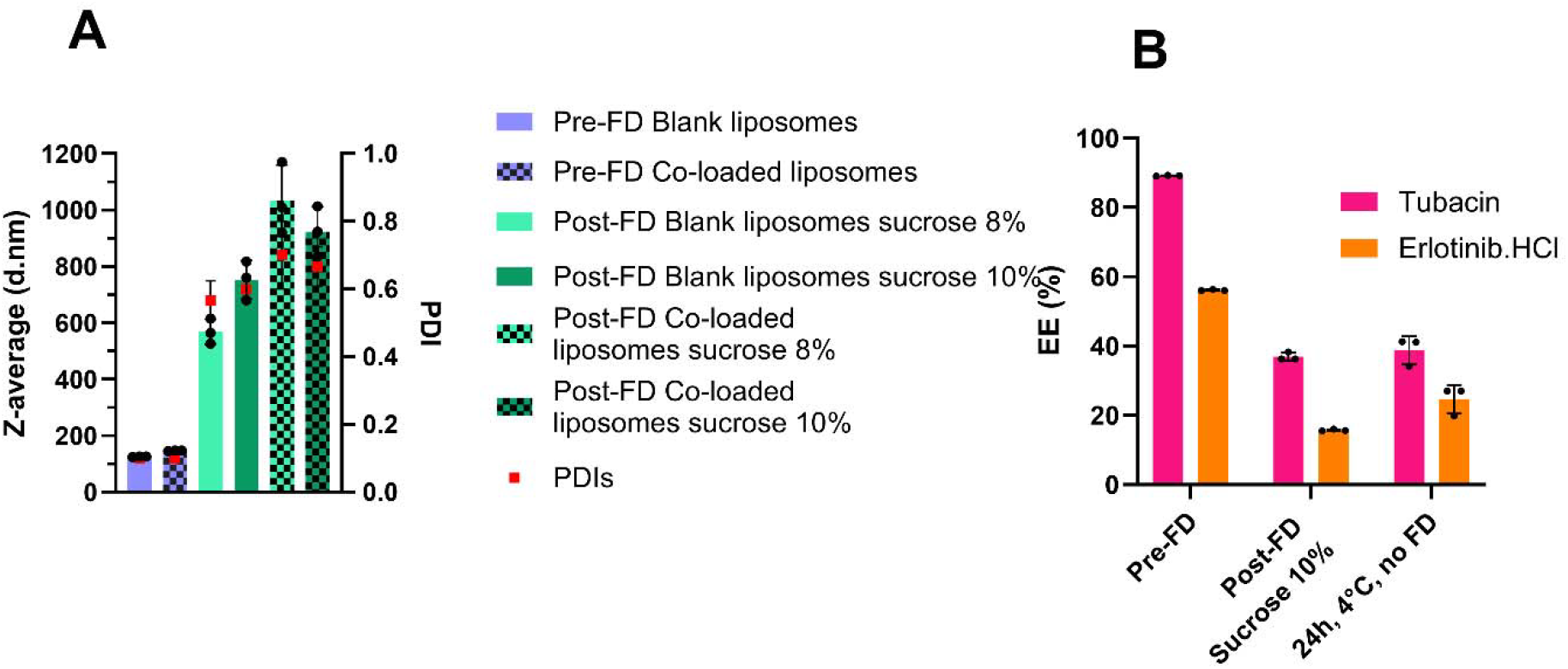
Z-average and PDIs of liposomes pre- and post-FD with the Christ^®^ freeze-dryer. Liposomes were either blank or co-loaded with tubacin and erlotinib. Intra-liposomal content was AS 120 mM, (N=1).

**Figure 6.**
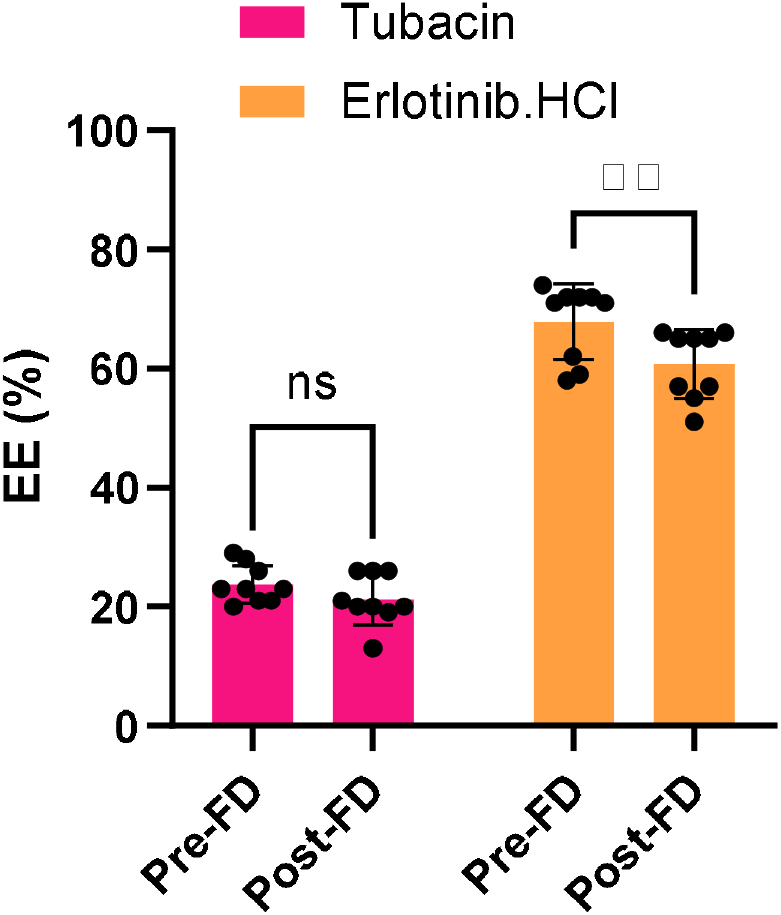
Encapsulation efficiencies of co-loaded liposomes pre- and post-FD with Christ^®^ freeze-dryer. AS concentration was 150 mM. N=3 ± SD. ** = p<0.01, ns = non-significance.

Formulations prepared with 150 mM AS demonstrated optimal characteristics, with Z-average diameters and PDIs closely matching pre-FD values. Based on these results, this molarity of AS was maintained as a standard concentration for the subsequent experiments. Liposomes were then lyophilized using the Telstar^®^ freeze-dryer, which enables precise control over freezing rates, primary drying parameters (temperature, pressure), as well as secondary drying steps. No differences were detected between pre-FD and post-FD sizes and PDIs using sucrose at 10% as a cryoprotectant (**Figure 7**). However, 8% led to significantly higher, although acceptable, sizes and PDIs.

**Figure 7.**
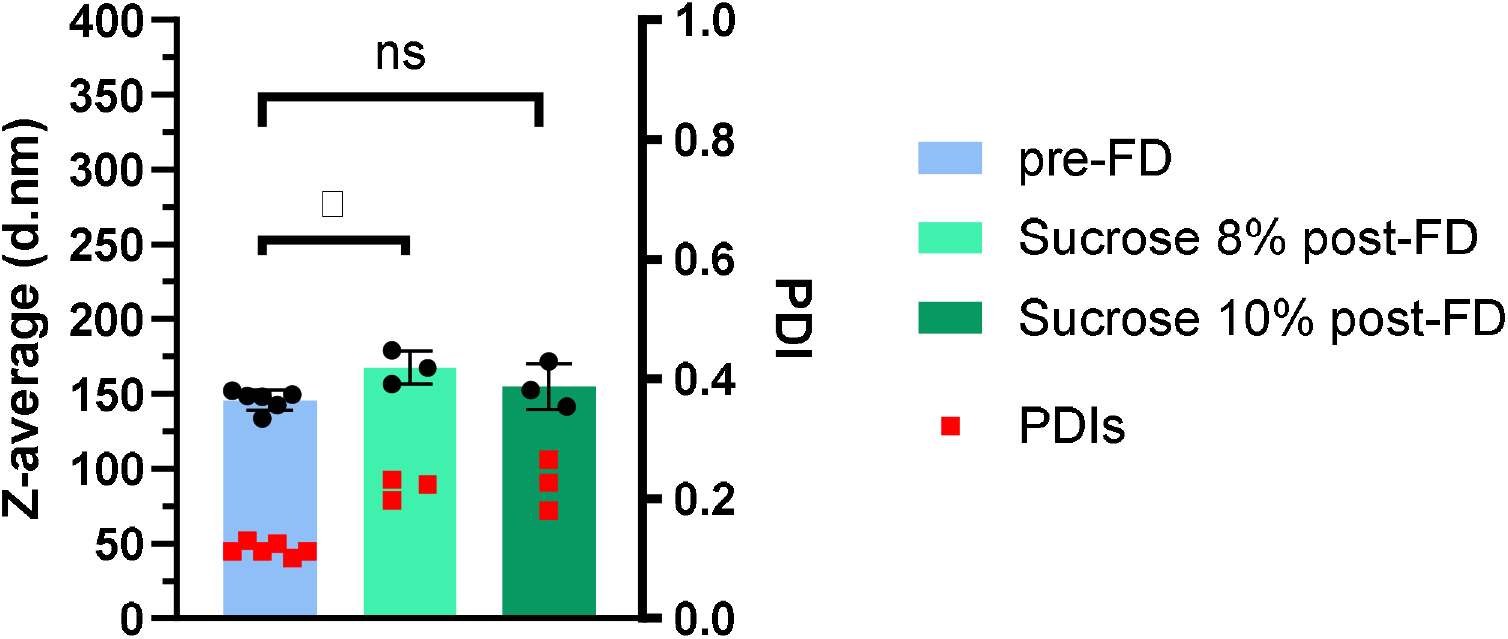
Z-average and PDIs of liposomes pre- and post-FD with different concentrations of sucrose. Freeze-drying performed with Telstar^®^. * = p<0.05, ns = non-significance. N=3, n=1 (post-FD) or n=3 ± SD (Pre-FD).

We further investigated the effect of intra-liposomal cryoprotectant incorporation by embedding sucrose during liposome synthesis at three concentrations (4.5%, 8% or 10%, respectively). Liposome sizes before FD were similar (with mean diameters of approx. 150 nm) with a polydispersity below 0.1. After FD, particle sizes did not significantly increase except for sucrose at 8%. Size was homogenously maintained across all groups (PDI below 0.3) after FD (**Figure 8**).

**Figure 8.**
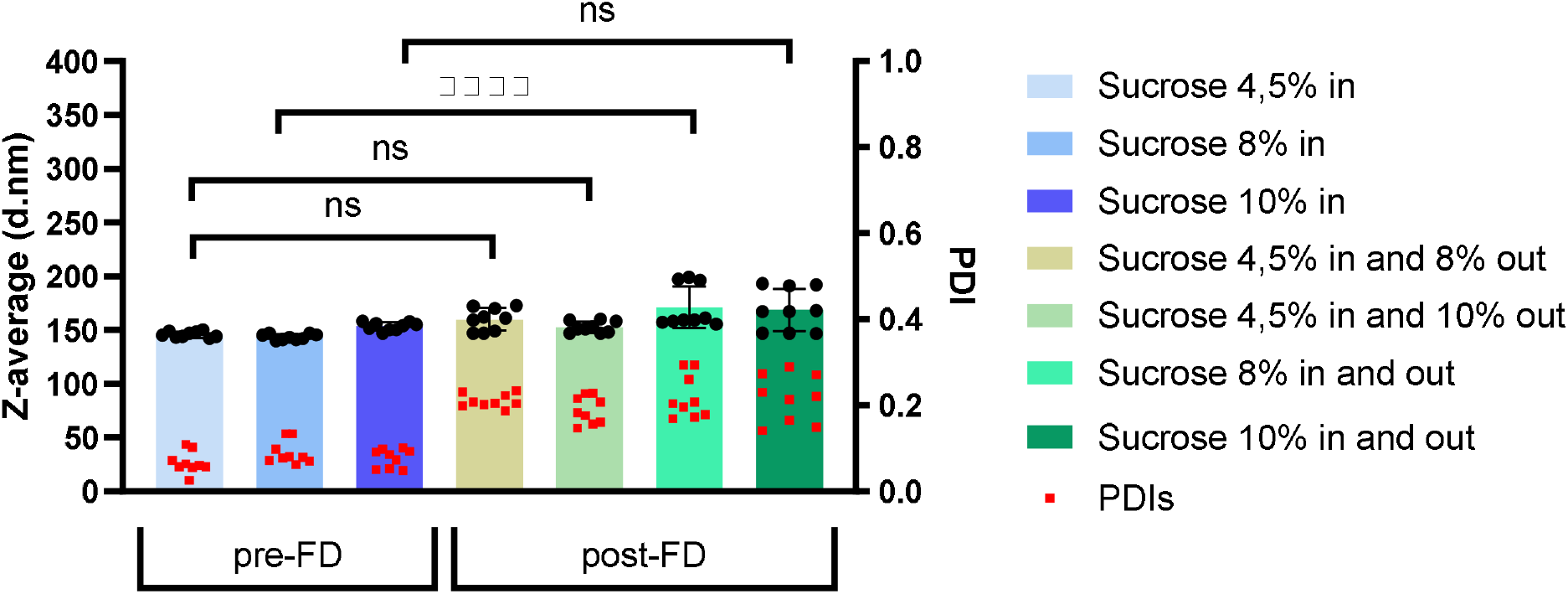
Z-average and PDIs of liposomes pre- and post-FD with different concentrations of sucrose in and out. Freeze-drying performed with Telstar^®^. N=3 ± SD. **** = p<0.0001, ns = non-significance.

The co-encapsulated liposomal formulation containing 10% sucrose exhibited excellent stability during freeze drying, with only approximately 10% API loss for both drugs during the freeze-drying process (**Figure 9A**). Dynamic light scattering (DLS) analysis in batch mode showed minimal changes in post-lyophilization, with a size increase of 20 nm while PDIs rose from 0.1 to 0.25 (**Figure 9B**). High-resolution AF4-MALS-DLS analysis confirmed exceptional preservation of particle properties identical 45-minute elution times pre- and post-lyophilization (**Figure 9C**). Monodisperse populations were maintained, as evidenced by a single monodisperse peak (**Figure 9D**), and unaltered conformation, as shown with unchanged shape factors constant at 0.9, **Figure 9E** and **9F**.

**Figure 9.**
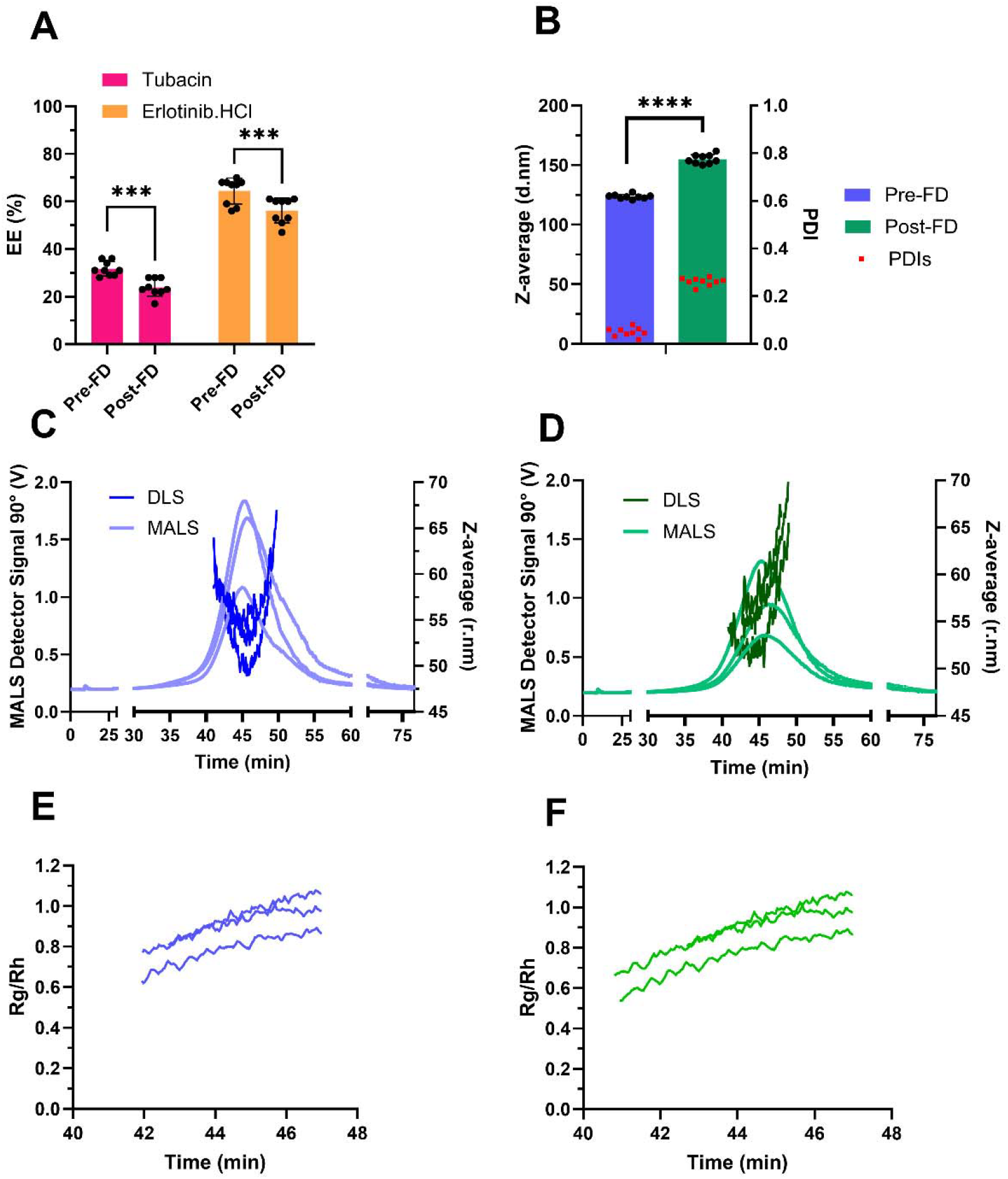
Characterization of co-loaded liposomes pre- and post-FD with the Telstar^®^. Encapsulation efficiency and drug concentrations in µM (**A**) and sizes with PDIs identified by DLS in batch-mode (**B**). AF4-DLS-MALS analysis of liposome pre-(**C**) and post-FD (**D**). Rg/Rh ratio or shape factor pre-(**E**) and post-FD (**F**). *** = p<0.001, **** = p<0.0001, ns = non-significance. N=3 ± SD.

## Discussion

The co-encapsulation of multiple therapeutic agents represents a promising strategy to mitigate drug losses during storage and administration while maintaining a synergistic ratio [6]. However, this strategy presents significant formulation challenges, particularly in achieving optimal drug loading while maintaining system stability. In this study, we developed and characterized liposomes co-encapsulating tubacin and erlotinib.HCl with particular focus on their storage stability at 4°C, physicochemical changes during the lyophilization process, as well as drug-loading preservation efficiency pre- and post-FD processing.

Tubacin is a highly hydrophobic compound and well-suited for passive loading into liposomes. Erlotinib.HCl is also a hydrophobic, weak-base API and is, therefore, charged positively in acid solutions. This feature makes it a promising candidate for remote loading into liposomes [21]. Our results demonstrated that lowering the internal liposomal pH resulted in increased encapsulation efficiency of erlotinib.HCl. The lowest pH tested (pH 3) was maintained for subsequent experiments. This observation suggests the charged nature of erlotinib.HCl may enhance its aqueous solubility and, therefore, improving its encapsulation efficiency. In the next step, we investigated various DSPC:Chol molar ratios, as cholesterol is known to play an important role in the reduction of phospholipid bilayer permeability and in increased API retention [22]. The DSPC:Chol ratio of 1:3.16 led to the highest erlotinib.HCl encapsulation. Conversely, tubacin encapsulation efficiency improved with lower cholesterol content, leading to enhanced encapsulation efficiency. This aligns with existing literature, where cholesterol is stated to compete for space within the bilayer with hydrophobic APIs [23].

Notably, all formulations, regardless of internal pH or DSPC:Chol molar ratios, exhibited high drug losses during storage at 4°C. This emphasized the necessity for an alternative stabilization strategy, leading to the development of the freeze-drying process as an effective solution. Freeze-drying is a process often chosen to mitigate stability challenges during liposome transport and storage. However, it is a process that induces a significant level of stress on the phospholipid bilayers. Freeze-drying of lipid-based nanoparticles is usually performed in three main steps, i.e., freezing, primary drying, and secondary drying [24]. During the freezing step, the formation of ice crystals can disrupt the liposomal membranes [25]. The primary drying phase involves sublimation, i.e., the withdrawal of water molecules from the formulation. This process can cause a tighter arrangement of the phospholipids within the bilayers, increasing van der Waals interactions. This phenomenon, in turn, can result in the fusion and aggregation of individual liposomes, leading to an increase in particle sizes [19]. Finally, the secondary drying phase is a critical step for the removal of the residual water content from the matrix [26]. This step is important as residual water content can decrease the glass transition temperature (Tg), a key parameter for ensuring the stabilization of liposomes [16]. To mitigate this, several types of cryoprotectants can be employed during the freeze-drying process of liposomes. Disaccharides are particularly effective, as they assure an optimal balance of sufficiently small size, reducing steric disturbance of the lipid bilayer and favoring interaction with phospholipids. They also have a high glass Tg, allowing them to remain in a glassy state for the longest time possible, leading to better stability of freeze-dried products [14, 27].

In this study, we tested the suitability of two disaccharides, i.e., sucrose and trehalose, at several concentrations. A total disaccharide concentration exceeding 2.5% (w/v) was required to achieve an effective cryoprotection [12]. Freeze-drying with trehalose resulted in a 4-fold increase in particle size, whereas the use of sucrose led to only a 2-fold size increase of approximately 150 nm. Although sucrose-containing formulations exhibited a modest size increase post-FD, the final particle size remained within the optimal range suitable for exploiting the enhanced permeability and retention (EPR) effect [28]. Furthermore, we observed a weak correlation between disaccharide concentrations with sucrose 6%, 8%, and 10%, leading to size expansion post-FD of 161 ± 1, 138 ± 1, 127 ± 1 nm, respectively. This concentration-dependent stabilization effect likely reflects the presence of more disaccharide units providing better protection and liposome integrity during the freezing and drying processes.

When selecting and evaluating the efficiency of a cryoprotectant for lyophilization process, the key parameter is the amorphous solid glass transition temperature (Tg), particularly the glass transition temperature of the maximally freeze-concentrated solution (Tg’). The Tg should exceed storage temperature by at least 10°C [29]. Trehalose demonstrates superior (Tg=117°C) compared to sucrose (Tg=77°C); however, both disaccharides exceed required storage temperature (2-8 °C) [27, 30]. Below the Tg’, the residual water in the freeze-dried product transitions to a glass state with minimal mobility, providing optimal protection during the FD process. The impact of each disaccharide’s Tg’ is minimal as they share similar Tg’ of −32°C and −30°C for sucrose and trehalose, respectively [26]. Trehalose was a superior cryoprotectant, although it was suggested to have a partial crystallization during the freeze-drying process [31, 32]. Additionally, its aqueous solubility was reduced compared to sucrose at temperatures below 60°C [33]. Roy *et al*. [34] observed through molecular dynamics simulations that sucrose formed more than 10% of hydrogen bonds with the polar heads of phospholipids than trehalose. The water replacement theory suggests that the formation of hydrogen bonds with phospholipids is essential for freeze-drying protection. This could explain why better results were obtained with sucrose compared to trehalose.

Previous studies have established that cryoprotectants such as carbohydrates and polymers cannnot permeate intact liposomal membrane [35]. Therefore, we studied the incorporation of cryoprotectants intra-liposomally for optimal protection as suggested by the literature [26, 36]. Inventors of Vyxeos^®^ lyophilized liposomes suggest a minimal cryoprotectant concentration of 125 mM for optimal stabilization [37]. Therefore, we added at least 4.5% of sucrose, corresponding to a concentration of 130 mM. Extra-liposomal cryoprotectant concentration was kept at either 8% or 10%. No statistically significant differences were observed among the various experimental conditions. The particle sizes measured before and after freeze-drying were comparable for sucrose concentrations of 4.5% and 10%. Our analysis revealed a significant divergence in size in liposomes containing 8% intra-liposomal sucrose compared to sucrose present both internally and externally post-FD. However, these results showed no significant variation in liposome size was detected when compared to previous results, where sucrose was exclusively present in the extra-liposomal medium. Based on these results, the subsequent phase of the study was conducted with the use of an external sucrose concentration of 10%.

Initial formulations were prepared using 150 mM ammonium sulfate as the intra-liposomal buffer for erlotinib.HCl remote loading. Osmolarity measurements indicated that blank liposomes combined with cryoprotectant exhibited approximately twice the osmolarity of the 150 mM AS buffer, potentially leading to a drug loss during encapsulation. To mitigate this, the formulations were adjusted to the concentration of 300 mM AS to match the osmolarity of liposomes combined with the cryoprotectant. However, these formulations demonstrated instability and precipitated shortly after preparation. This instability was attributed to the excessive permeation of ammonium ions (NH_4_^+^), which may cause pore formation in the bilayer and further lipid precipitation [38]. We subsequently evaluated 120 mM AS to match PBS osmolarity. No precipitation was observed, however, post-FD, reconstituted liposomes featured sizes above 600 nm, high polydispersity (PDIs above 0.5), and substantial drug loss. The molarity of ammonium sulfate used for API remote encapsulation into liposomes varies in the literature. Chen *et al*. [39] used 200 mM AS, while Jun-Jen *et al*. used 250 mM AS with an intra-liposomal osmolarity of 520 mOs, which was paired with NaCl 0.9% as the external buffer (286 mOs) [40]. Zhigaltsev *et al*. prepared liposomes using 120 mM AS [41]. Another study correlated the ammonium sulfate molarity with doxorubicin sulfate nanorod crystals and determined a baseline molarity of 200 mM to achieve a stable nanocrystallization of doxorubicin [42]. It appears that the molarity of ammonium sulfate is, therefore formulation specific. Based on preliminary optimization, 150 mM AS was selected as the standard condition for subsequent evaluations in this study.

Subsequently, two freeze-drying protocols were followed and compared. For the FD process conducted with a Christ^®^ Alpha 2-4 LSC freeze-dryer, the freezing step was performed at −80°C in an external freezer, followed by a primary drying at −40°C for 24 hours within the freeze-dryer. Although this initial method yielded satisfactory API retention and particle size, its utility was limited by the absence of a secondary drying phase and a freezing step. In contrast, the Telstar^®^ lyoBeta mini freeze-dryer enabled precise control over the different steps in the freeze-drying process. Indeed, the freezing process (−40°C for 3 hours), primary drying (−40°C to −10°C for over 33 hours), and secondary drying (22°C for 4 hours) were performed within a single apparatus. This integrated system provided real-time monitoring of both temperature and pressure, ensuring the optimal process.

Comparative analysis of cooling rates revealed rapid (1.33°C/min) cooling using the Christ^®^ freeze-dryer, while the cooling rate for the Telstar^®^ was slower (0.22°C/min). Slow cooling rates, such as 0.5°C/min are favorable to avoid immediate ice formation, which would disrupt the integrity of the phospholipid bilayers [26]. As previously reported, primary drying is usually performed at low pressures between 0.03 and 1 mbar with temperatures ranging between −50°C and −60°C [43]. The primary drying in this study was done at −40°C with a pressure of 0.3 mbar, reflecting the operational limit of the Telstar^®^ system. During the freezing and primary drying phases, temperatures were maintained at −40°C (below the Tg’) to ensure prevention of lyophilized cake collapse [44] and preservation of liposome structural integrity.

The optimized liposomal formulation co-encapsulated with tubacin and erlotinib.HCl lyophilized using the Telstar^®^ system with sucrose at 10%. There was a significant API loss of 10% during this process. Using DLS in batch mode, we observed an increase in particle size. However, this size change was not found in DLS flow mode, as the elution times of the particles remained similar during pre- and post-FD process. This observed size increase in batch mode DLS was insufficient to alter the elution profile of the particles during the analysis. AF4-MALS-DLS results showed a single peak, indicating preserved monodisperse size distribution. The AF4-DLS system provided a robust, high-resolution size distribution and separation [45]. Further analysis through the calculated shape factor using the gyration radius (Rg) divided by the hydrodynamic radius (Rh) from DLS. The obtained ratio of Rg/Rh at approximately 0.9 suggested that particles are spherical and retain this shape after the freeze-drying process [46].

## Conclusions

In this study, we established an optimized process for co-encapsulation of tubacin and erlotinib.HCl into liposomal formulations while systematically evaluating formulation parameters (internal pH, DSPC-to-cholesterol ratios, or single-*vs*. co-encapsulation), stabilization challenges (drug loss during storage at 4°C), and product stabilization by freeze-drying. Our results show that sucrose outperformed trehalose as a cryoprotectant, exhibiting better stabilization of particle sizes and PDI. This is likely due to sucrose’s higher capacity to form hydrogen bonds with phospholipids. The Telstar^®^ freeze-dryer’s precise temperature control over freezing and drying phases, enabled maintenance of critical process conditions below the Tg’ ensuring optimal liposome protection. Liposomes freeze-dried with 10% sucrose showed minimal drug loss but maintained optimal physicochemical characteristics (particle size, and spherical morphology to exploit the EPR-mediated tumor effect. While the freeze-drying process showed encouraging results, further improvements are required in the future to minimize drug loss and enhance size stability. Further studies should focus on a refinement of process optimization with the assessment of residual moisture content and testing of different drying phase parameters for different durations in order to achieve an improved formulation suitable for further clinical translation and potential therapeutic applications.

## Competing interests

The authors have declared no competing interest.

## References

1. Chou, T.C., Drug combination studies and their synergy quantification using the Chou-Talalay method. Cancer Res, 2010. 70(2): p. 440–6 10.1158/0008-5472.CAN-09-1947.

2. Mayer, L.D., et al., Ratiometric dosing of anticancer drug combinations: controlling drug ratios after systemic administration regulates therapeutic activity in tumor-bearing mice. Mol Cancer Ther, 2006. 5(7): p. 1854–63 10.1158/1535-7163.MCT-06-0118.

3. Weiss, A., et al., Identification of a Synergistic Multi-Drug Combination Active in Cancer Cells via the Prevention of Spindle Pole Clustering. Cancers (Basel), 2019. 11(10): p. 1612–34 10.3390/cancers11101612.

4. Kumar, V. and N. Dogra, A Comprehensive Review on Deep Synergistic Drug Prediction Techniques for Cancer. Archives of Computational Methods in Engineering, 2021. 29(3): p. 1443–1461 10.1007/s11831-021-09617-3.

5. Kraft, J.C., et al., Emerging research and clinical development trends of liposome and lipid nanoparticle drug delivery systems. J Pharm Sci, 2014. 103(1): p. 29–52 10.1002/jps.23773.

6. Zununi Vahed, S., et al., Liposome-based drug co-delivery systems in cancer cells. Mater Sci Eng C Mater Biol Appl, 2017. 71: p. 1327–1341 10.1016/j.msec.2016.11.073.

7. Kemp, J.A., et al., “Combo” nanomedicine: Co-delivery of multi-modal therapeutics for efficient, targeted, and safe cancer therapy. Adv Drug Deliv Rev, 2016. 98: p. 3–18 10.1016/j.addr.2015.10.019.

8. Zhang, R.X., et al., Nanomedicine of synergistic drug combinations for cancer therapy - Strategies and perspectives. J Control Release, 2016. 240: p. 489–503 10.1016/j.jconrel.2016.06.012.

9. Qu, M.H., et al., Liposome-based co-delivery of siRNA and docetaxel for the synergistic treatment of lung cancer. Int J Pharm, 2014. 474(1-2): p. 112–22 10.1016/j.ijpharm.2014.08.019.

10. Jia, L., et al., Characterization techniques: The stepping stone to liposome lyophilized product development. Int J Pharm, 2021. 601: p. 120519 10.1016/j.ijpharm.2021.120519.

11. Luo, W.C., et al., Impact of formulation on the quality and stability of freeze-dried nanoparticles. Eur J Pharm Biopharm, 2021. 169: p. 256–267 10.1016/j.ejpb.2021.10.014.

12. Mohammed, A.R., et al., Lyophilisation and sterilisation of liposomal vaccines to produce stable and sterile products. Methods, 2006. 40(1): p. 30–8 10.1016/j.ymeth.2006.05.025.

13. Wang, Y. and D.W. Grainger, Lyophilized liposome-based parenteral drug development: Reviewing complex product design strategies and current regulatory environments. Adv Drug Deliv Rev, 2019. 151-152: p. 56–71 10.1016/j.addr.2019.03.003.

14. Stark, B., G. Pabst, and R. Prassl, Long-term stability of sterically stabilized liposomes by freezing and freeze-drying: Effects of cryoprotectants on structure. Eur J Pharm Sci, 2010. 41(3-4): p. 546–55 10.1016/j.ejps.2010.08.010.

15. Crowe, J.H. and L.M. Crowe, Factors affecting the stability of dry liposomes. Biochim Biophys Acta, 1988. 939(2): p. 327–34 10.1016/0005-2736(88)90077-6.

16. Ingvarsson, P.T., et al., Stabilization of liposomes during drying. Expert Opin Drug Deliv, 2011. 8(3): p. 375–88 10.1517/17425247.2011.553219.

17. Crowe, J.H., S.B. Leslie, and L.M. Crowe, Is vitrification sufficient to preserve liposomes during freeze-drying? Cryobiology, 1994. 31(4): p. 355–66 10.1006/cryo.1994.1043.

18. Koster, K.L., et al., Interactions between soluble sugars and POPC (1-palmitoyl-2-oleoylphosphatidylcholine) during dehydration: vitrification of sugars alters the phase behavior of the phospholipid. Biochim Biophys Acta, 1994. 1193(1): p. 143–50 10.1016/0005-2736(94)90343-3.

19. Franze, S., et al., Lyophilization of Liposomal Formulations: Still Necessary, Still Challenging. Pharmaceutics, 2018. 10(3): p. 139–62 10.3390/pharmaceutics10030139.

20. Guimaraes, D., et al., Protective Effect of Saccharides on Freeze-Dried Liposomes Encapsulating Drugs. Front Bioeng Biotechnol, 2019. 7: p. 424–32 10.3389/fbioe.2019.00424.

21. Cern, A., et al., New drug candidates for liposomal delivery identified by computer modeling of liposomes’ remote loading and leakage. J Control Release, 2017. 252: p. 18–27 10.1016/j.jconrel.2017.02.015.

22. Briuglia, M.L., et al., Influence of cholesterol on liposome stability and on in vitro drug release. Drug Deliv Transl Res, 2015. 5(3): p. 231–42 10.1007/s13346-015-0220-8.

23. Ali, M.H., et al., Solubilisation of drugs within liposomal bilayers: alternatives to cholesterol as a membrane stabilising agent. J Pharm Pharmacol, 2010. 62(11): p. 1646–55 10.1111/j.2042-7158.2010.01090.x.

24. Wang, L., et al., Cryoprotectant choice and analyses of freeze-drying drug suspension of nanoparticles with functional stabilisers. J Microencapsul, 2018. 35(3): p. 241–248 10.1080/02652048.2018.1462416.

25. Liu, B., et al., Potential of trehalose and sucrose-raffinose combination instead of sucrose applied to the cytarabine/daunorubicin co-loaded lyophilized liposome: Investigation on the protective effect and mechanism of various lyoprotectants. J. drug del. sci. tech, 2024. 92: p. 105360–9 10.1016/j.jddst.2024.105360.

26. van Winden, E.C., Freeze-drying of liposomes: theory and practice. Methods Enzymol, 2003. 367: p. 99–110 10.1016/S0076-6879(03)67008-4.

27. Tonnis, W.F., et al., Size and molecular flexibility of sugars determine the storage stability of freeze-dried proteins. Mol Pharm, 2015. 12(3): p. 684–94 10.1021/mp500423z.

28. Wu, J., The Enhanced Permeability and Retention (EPR) Effect: The Significance of the Concept and Methods to Enhance Its Application. J Pers Med, 2021. 11(8) 10.3390/jpm11080771.

29. Mensink, M.A., et al., How sugars protect proteins in the solid state and during drying (review): Mechanisms of stabilization in relation to stress conditions. Eur J Pharm Biopharm, 2017. 114: p. 288–295 10.1016/j.ejpb.2017.01.024.

30. Guarro, M., et al., Efficient extracellular vesicles freeze-dry method for direct formulations preparation and use. Colloids Surf B Biointerfaces, 2022. 218: p. 112745 10.1016/j.colsurfb.2022.112745.

31. Kannan, V., et al., Effect of sucrose as a lyoprotectant on the integrity of paclitaxel-loaded liposomes during lyophilization. J Liposome Res, 2015. 25(4): p. 270–8 10.3109/08982104.2014.992023.

32. Sundaramurthi, P. and R. Suryanarayanan, Trehalose Crystallization During Freeze-Drying: Implications On Lyoprotection. J. Phys. Chem. Lett., 2009. 1(2): p. 510–514 10.1021/jz900338m.

33. Sundaramurthi, P., T.W. Patapoff, and R. Suryanarayanan, Crystallization of trehalose in frozen solutions and its phase behavior during drying. Pharm Res, 2010. 27(11): p. 2374–83 10.1007/s11095-010-0243-2.

34. Roy, A., et al., A Comparative Study of the Influence of Sugars Sucrose, Trehalose, and Maltose on the Hydration and Diffusion of DMPC Lipid Bilayer at Complete Hydration: Investigation of Structural and Spectroscopic Aspect of Lipid-Sugar Interaction. Langmuir, 2016. 32(20): p. 5124–34 10.1021/acs.langmuir.6b01115.

35. Boafo, G.F., et al., The Role of Cryoprotective Agents in Liposome Stabilization and Preservation. Int J Mol Sci, 2022. 23(20) 10.3390/ijms232012487.

36. Arshinova, O.Y., et al., Lyophilization of liposomal drug forms (Review). Pharm. Chem. J., 2012. 46(4): p. 228–233 10.1007/s11094-012-0768-2.

37. Donna Cabral-Lilly, L.M., Paul Tardi, David Watkins, Yi Zeng, Lyophilized liposomes. 2012, Jazz Pharmaceuticals Research LLC.

38. Abraham, S.A., et al., An evaluation of transmembrane ion gradient-mediated encapsulation of topotecan within liposomes. J Control Release, 2004. 96(3): p. 449–61 10.1016/j.jconrel.2004.02.017.

39. Chen, J., et al., Ammonium sulfate gradient loading of brucine into liposomes: effect of phospholipid composition on entrapment efficiency and physicochemical properties in vitro. Drug Development and Industrial Pharmacy, 2010. 36(3): p. 245–253 10.3109/03639040903099736.

40. Liu, J.-J., et al., Simple and efficient liposomal encapsulation of topotecan by ammonium sulfate gradient: stability, pharmacokinetic and therapeutic evaluation. Anti-Cancer Drugs, 2002. 13(7): p. 709–717.

41. Zhigaltsev, I.V., et al., Liposomes containing dopamine entrapped in response to transmembrane ammonium sulfate gradient as carrier system for dopamine delivery into the brain of parkinsonian mice. J Liposome Res, 2001. 11(1): p. 55–71 10.1081/LPR-100103170.

42. Wei, X., et al., Cardinal Role of Intraliposome Doxorubicin-Sulfate Nanorod Crystal in Doxil Properties and Performance. ACS Omega, 2018. 3(3): p. 2508–2517 10.1021/acsomega.7b01235.

43. Pardeshi, S.R., et al., Process development and quality attributes for the freeze-drying process in pharmaceuticals, biopharmaceuticals and nanomedicine delivery: a state-of-the-art review. Futur. J. Pharm. Sci., 2023. 9(1): p. 99–130 10.1186/s43094-023-00551-8.

44. Hussain, M.T., et al., Freeze-drying cycle optimization for the rapid preservation of protein-loaded liposomal formulations. Int J Pharm, 2020. 573: p. 118722 10.1016/j.ijpharm.2019.118722.

45. Clogston, J.D., The importance of nanoparticle physicochemical characterization for immunology research: What we learned and what we still need to understand. Adv Drug Deliv Rev, 2021. 176: p. 113897 10.1016/j.addr.2021.113897.

46. Hu, Y., R.M. Crist, and J.D. Clogston, The utility of asymmetric flow field-flow fractionation for preclinical characterization of nanomedicines. Anal Bioanal Chem, 2020. 412(2): p. 425–438 10.1007/s00216-019-02252-9.

